# Preparation and optimization of MEPEG-PLGA nanoparticles for gene delivery

**DOI:** 10.1101/624254

**Authors:** Leah Monash, John Smith

## Abstract

This paper demonstrates a method to prepare a cationic methoxy-terminated polyethylene glycol-polylactic acid polyglycolic acid block polymer (MePEG-PLGA) nanoparticle by a nanoparticle precipitation method was established. This study used single factor design and orthogonal experiment to select the optimal experimental scheme and examined the physical properties of the nanoparticles such as surface morphology, particle size distribution, zeta potential, DNA binding rate, and DNA protection ability. Results indicate that the optimal size of the prepared nanoparticles was 89.7 nm and the surface potential was 28.3 mV. The nanoparticles were scattered under transmission electron microscope, the size was uniform, the surface was smooth, and the distribution was spherical. The DNA binding rate was 80.2 %, and can well protect the contained genes from nuclease degradation. Conclusion The cationic nanoparticles prepared by nanoparticle precipitation method are expected to be highly efficient gene carriers. The preparation of cationic methoxy-terminated polyethylene glycol-polylactic acid polyglycolic acid block polymer (MePEG-PLGA) by nanoparticle precipitation method. Nanoparticle method. Method This study used single factor design and orthogonal experiment to select the optimal experimental scheme, and examined the physical properties of the nanoparticles such as surface morphology, particle size distribution, Zeta potential, DNA binding rate, and DNA protection ability. Results The optimal size of the prepared nanoparticles was 89.7 nm and the surface potential was 28.3 mV. The nanoparticles were scattered under transmission electron microscope, the size was uniform, the surface was smooth, and the distribution was spherical. The DNA binding rate was 80.2 %, and can well protect the contained genes from nuclease degradation. Conclusion The cationic nanoparticles prepared by nanoparticle precipitation are expected to be efficient gene carriers.

## Introduction

Studies have found that neurotrophic factors (NTFs) drugs can promote and maintain neuronal growth, survival, differentiation and executive function, and can effectively improve Alzheimer’s disease (AD) symptoms. However, current NTFs drugs have not been used in clinical treatment because NTFs are difficult to pass through the blood-brain barrier, and frequent intracerebral injections have the potential to cause new damage or infection in the brain. Transgenic therapy has received more and more attention, but the carrier problem of gene therapy, as well as carrier-related immune response, cytotoxicity and safety issues are the bottlenecks limiting the development of gene therapy and clinical application [1]. Gene vectors must be characterized by safety, stability, and high transfection efficiency. Non-viral vectors have no immunogenicity and no genotoxicity compared with viral vectors [2]. Nanocarriers have attracted more and more research and attention due to their unique advantages such as structural designability, particle size controllability, sustained release, and smaller particle size [2–3]. Studies have shown that when the particle size of nanoparticles is less than 100 nm, the transfection efficiency is 27 times higher than that of nanoparticles with 100 nm [4]. Poly(D,L-lactide-co-glycolide, PLGA) is a biodegradable, biocompatible material approved by the US FDA for use in humans [5]. It is widely used in tissue engineering and drug carrier research. It has the dual role of scaffold and sustained release, and has potential value as a controlled release system for plasmid DNA. However, the hydrophobicity of PLGA limits its advantages. Polyethylene glycol (PEG) is a non-ionic hydrophilic segment. It is the most commonly used hydrophilic modification molecule. It has excellent biocompatibility, strong hydrophilicity, high cell permeability, and is nontoxic in the body. Immunogenicity [6] has been approved by the FDA for use in humans. The introduction of a hydrophilic PEG segment into the hydrophobic PLGA molecule improves the hydrophilicity and flexibility of the polymer, enhances particle stability and enzyme activity, and improves interaction with cellular components. At the same time, the hydrophilic microenvironment formed by PEG can improve the efficiency of DNA loading and is beneficial to the maintenance of gene drug activity. Cetyl trimethyl ammonium bromide (CTAB) is a cationic surfactant with good stability and biodegradability. Using CTAB to trim MePEG-PLGA nanoparticles, the nanoparticle surface can be positively charged to efficiently bind negatively charged DNA [7–8]. There are many methods for preparing polymer nanoparticles. In this study, nanoparticle precipitation method was selected [9–10]. This method does not require external forces such as ultrasound and high-speed shear, nor does it introduce high-temperature, toxic organic solvents, etc., which is beneficial to maintain the biological activity of bio-sensitive macromolecules [11]. Cationic MePEG-PLGA nanoparticles have the following advantages: biodegradable materials, intermediate metabolites (lactic acid and glycolic acid) and final degradation products (CO2 and H2O) are not toxic to cells; no immunogenicity, no immune response Because the surface is positively charged, it can adsorb and concentrate DNA and protect the gene from degradation; the larger surface area is good for DNA adsorption and depolymerization; the smaller particle size is expected to enter the cell by endocytosis. The cation ion MePEG-PLGA nanoparticles become a highly efficient and non-toxic novel gene vector. The purpose of this study was to investigate the effects of different factors on the particle size of nanoparticles and to select the optimal protocol for the next step of transfecting cells.

## Materials and Method

Agarose (BIOWEST, Spain); MePEG-PLGA (Shandong Biotech Co., Ltd.); CTAB (Amersco, USA); DNase I (Biosharp, Japan); Constant Temperature Air Oscillation Bath (Harbin Dongming Medical); Ultrasonic Cell Crusher (Scientz-IID, Ningbo Xinzhi); Magnetic Stirrer (IKA, Germany); Low Temperature High Speed Centrifuge (Sigma, Germany); Transmission Electron Microscope (JEM-2010, Japan); Particle Size Analyzer (Zetasizer NanoZS90, Malvern, UK Company); UV spectrophotometer (HITACHI, Japan); micropipette (eppendorf, Germany).

LGA nanoparticle (hereinafter referred to as MePEG-PLGA-NPs). The operation is as follows: Pre-weigh the prescribed amount of MePEG-PLGA, ultrasonically dissolved in a certain volume of acetone, and the polymer solution is added to a 20 mL concentration of CTAB emulsifier under magnetic stirring with a 9-gauge needle at a certain flow rate. In an aqueous solution, continue stirring at room temperature overnight, completely evaporate the organic solvent, 15 000 r / min (about 20 000 g), centrifuge at 20 ° C for 20 min, wash with double distilled water to remove excess emulsifier, and then suspend the precipitate in double steaming. In water, centrifuge at 5 000 r / min (about 2 200 g) for 10 min, discard the large particles in the precipitate, and the supernatant is MePEG-PLGA-NPs suspension.

Single factor study According to the previous experimental results of the research group, factors that may affect the size of the nanoparticles were selected for single factor analysis: MePEG-PLGA concentration (A, mg / mL, in the oil phase), CTAB concentration (B, w / v)), injection rate (C, mL / min), magnetic stirring speed (D, r / min) and volume (E, mL), 5 factors for each factor (Table 1).

Select MePEG-PLGA concentration (A, mg / mL, oil phase), CTAB concentration (B, w / v), injection speed (C, mL / min) and magnetic stirring speed (D, r / min) 4 factors Each factor was taken at 3 levels according to the Lg (34) orthogonal experimental design table design experiment. The nanoparticle size was used as the evaluation index to comprehensively evaluate each prescription. The orthogonal design factor level table is shown in Table 2.

Preparation of genetically-loaded nanoparticles. Accurately measure the appropriate amount of DNA solution and MePEG-PLGA nanoparticles to make the DNA to nanoparticle mass ratio 1:100, vortex for 5 s, mix well, incubate for 20 min at room temperature, and obtain the cationic gene-loaded MePEG-PLGA nanoparticles (hereinafter referred to as DNA-MePEG-PLGA-NPs) were assayed for DNA binding by 0.8% agarose gel electrophoresis.

Determination of DNA binding rate Precision measurement of DNAMePEG-PLGA-NPs 200 μL, low temperature high-speed centrifugation (15 000 r / min, 20 000 g, 4 °C) for 20 min, then extract the supernatant 100 μL, determined by UV spectrophotometry The concentration of free DNA in the clear solution (with blank nanoparticle supernatant minus the blank), according to the formula “DNA binding rate = (total DNA amount – free D

The morphology of nanoparticles was observed by transmission electron microscopy. Specific methods: MePEG-PLGA-NPs and DNA96 were obtained separately. Anatomy of the study 2011, Vol. 33 No. 2 Anat Res, 2011, Vol. 33, No. 2 MePEG-PLGA-NPs suspension is a little, dripped onto the electron microscope copper net with carbon film, left to stand for 40 s, the remaining suspension is removed by filter paper, and the mass concentration is 2% phosphotungstic acid (dissolved) In PBS, the cells were negatively stained for 30 s. After drying naturally, the morphology of the nanoparticles was observed under a transmission electron microscope and photographed. The nanoparticle size and Zeta potential analyzer were used to determine the particle size distribution and Zeta potential of the nanoparticles: take appropriate amount of the nanoparticle suspension, and after appropriate dilution, measure with Zetasizer Nano-ZS90 particle size analyzer, dynamic light scattering software data processing, record average Particle size, polydispersity index and zeta potential.

Protecting plasmid DNA from nuclease degradation is one of the essential properties of gene vectors for efficient in vivo and in vitro gene delivery. Accurately measure 50 μL of DNA-MePEG-PLGA-NPs (containing plasmid DNA 1 μg), add 5.5 μL of DNase I Buffer (50 mmol / L Tris-HCl, 10 mmol / L MgCl2, pH 7.6), mix Evenly, add 2 μL (0.1 U / μg DNA) of 0.05 U / μL DNase I in 0.9% NaCl solution and mix. Another 1 μg of naked DNA was added, and 5 μL of DNase I Buffer and 0.05 μl / μL of DNase I 0.9 μ NaCl solution 2 μL (0.1 U / μg DNA) were added and mixed. The mixture was incubated on a 37 ° C air bath oven for 30 min (100 r / min), then 5 μL EDTA (0.5 mol / L, pH 8.0) was added for 15 min at room temperature to terminate the reaction. The group was added with a final concentration of 0.1% heparin [14–15] at 37 ° C in a water bath at a constant temperature for 3.5 h (100 r / min), replacing the DNA in the extraction system, and performing agarose gel simultaneously with the naked DNA. (0.8%) Electrophoresis block analysis (90 V, 25 min) was performed to analyze the protective ability of the nanoparticle vector for the contained gene.

## Results and Discussion

The results of single factor investigation showed that the particle size of nanoparticles increased with the increase of MePEG-PLGA concentration in organic solvents. However, when the amount of the polymer material is too small, the yield of the nanoparticles is low; when the amount is too high, the diameter of the nanoparticles is large, and agglomeration is prone to occur. CTAB is a cationic surfactant. Within a certain range, increasing the CTAB concentration can significantly reduce the particle size of the nanoparticles, and further increase its concentration, and the particle size of the nanoparticles will increase. The particle size of nano-particles increased with the increase of instillation rate, and the difference of nano-particle size was not significant when the instillation rate was 1 mL / min and 2.5 mL / min. The magnetic stirring speed has a certain influence on the particle size of the nanoparticles. The nanometer particle size decreases with the increase of the magnetic stirring speed. However, under the transmission electron microscope, it is found that the edge of the nanoparticle obtained when the stirring speed reaches 300 r / min or more Broken, incomplete form. The amount of acetone in the prescription significantly affects the particle size of the nanoparticles, but when the volume of acetone is increased from 5 mL to 7 mL, the particle size of the nanoparticles is not significantly reduced, and it takes longer to volatilize the organic solvent, so the volume of 5 mL is the most Excellent condition. Single-factor investigation The results of nano-particle size of each experimental group are shown in Table 3.

On the basis of single factor investigation, the nanoparticle formulation was optimized by orthogonal design to prepare cationic MePEG-PLGA-NPs with small particle size and moderate positive charge. According to the range R, the order of influence of the four factors on the particle size of the nanoparticles is A > D > C > B, in which the analysis results of each factor level are A: 3 > 2 > 1; B: 3 > 2 > small key Factors, but significantly affect the nanoparticle Zeta potential. In order to obtain the cations MePEG-PLGA-NPs with a small particle size and a high Zeta potential, the concentration of CTAB is chosen to be 0.5%, and the best position is A1B2C3D3. The results of the intersection are shown in Table 4. Three batches of samples were prepared according to the optimal prescription, and the average particle diameter and Zeta potential were measured respectively. The results showed that the optimal formulation optimized by orthogonal design had better formulation stability. The results are shown in Table 5.

The particle size distribution of the prepared nanoparticles is shown in Figure 1. The average particle size is 86.1 nm, the particle size distribution is uniform, the distribution index is 0.135, and the Zeta potential is 27.9 mV, indicating that the surface of the nanoparticles has a high positive The charge is expected to efficiently bind to the negatively charged plasmid DNA. The average particle size of the nanoparticles after adsorption of plasmid DNA was 103.3 nm, the distribution index was 0.222, and the Zeta potential was 14.1 mV. Photomicrographs of MePEGPLGA-NPs and DNA-MePEG-PLGA-NPs are shown in Figure 2. The nanoparticles are in the form of regular spheres or close to spheres.

**Figure 1.**
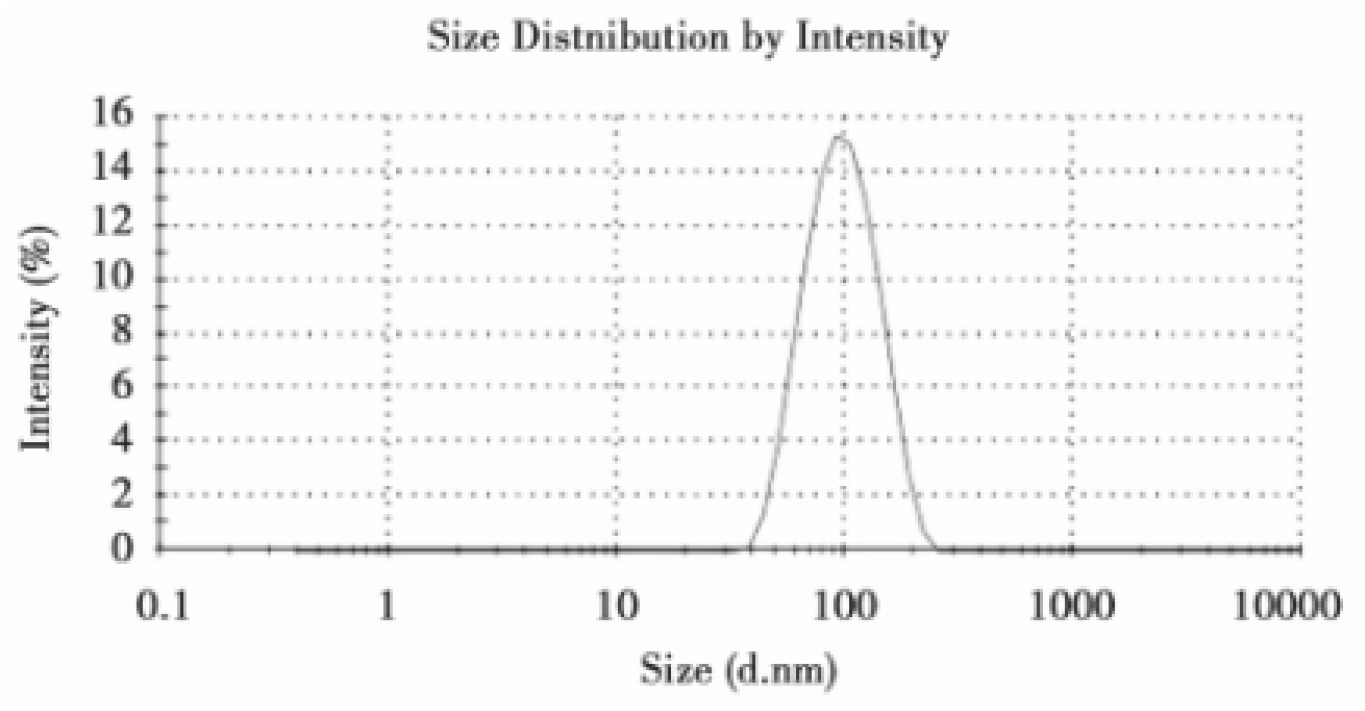
Size analysis for dynamic light scattering.

**Figure 2.**
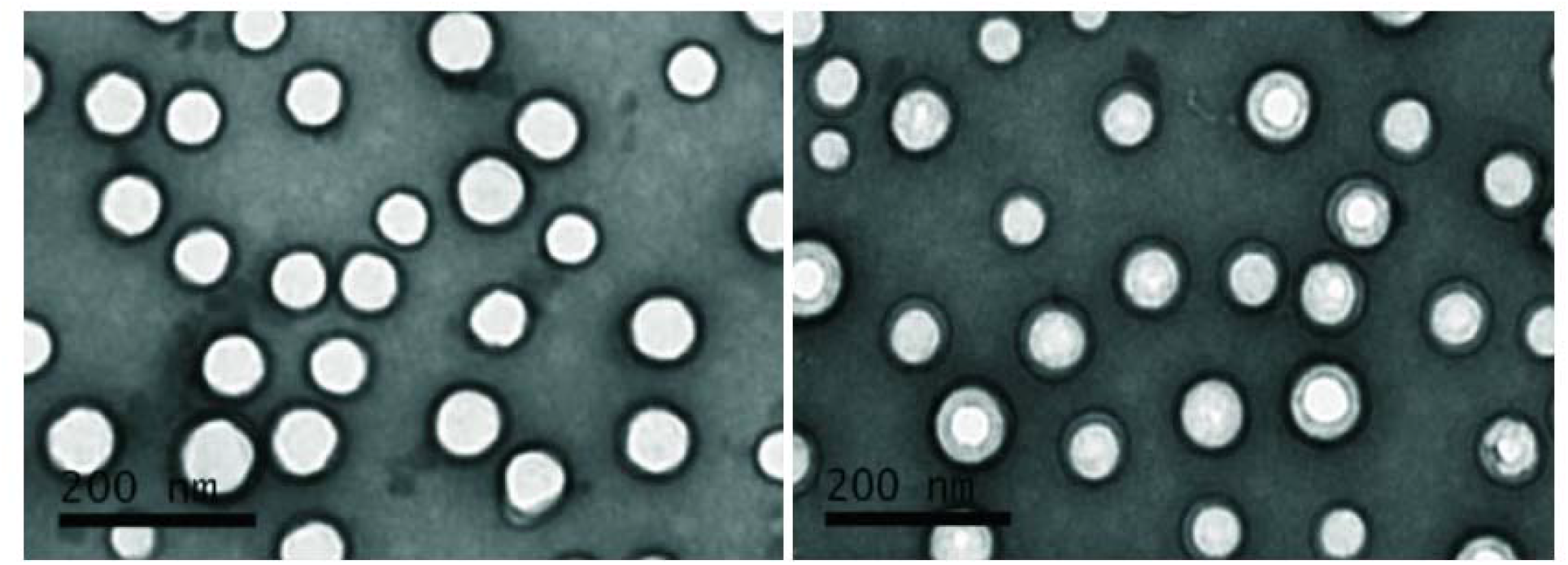
Nanoparticle morphology analysis by TEM

The DNA binding rate was determined by an ultraviolet spectrophotometer to be 80.0%. The electrophoresis results are shown in Figure 3. The plasmid DNA was bound to MePEG-PLGA-NPs and retained in the well. There was almost no free DNA in front, indicating that the DNA was completely integrated with MePEG-PLGA-NPs and formed a more stable DNA. MePEG-PLGA-NPs are consistent with DNA binding by UV spectrometry^1–11^.

**Figure 3 & 4.**
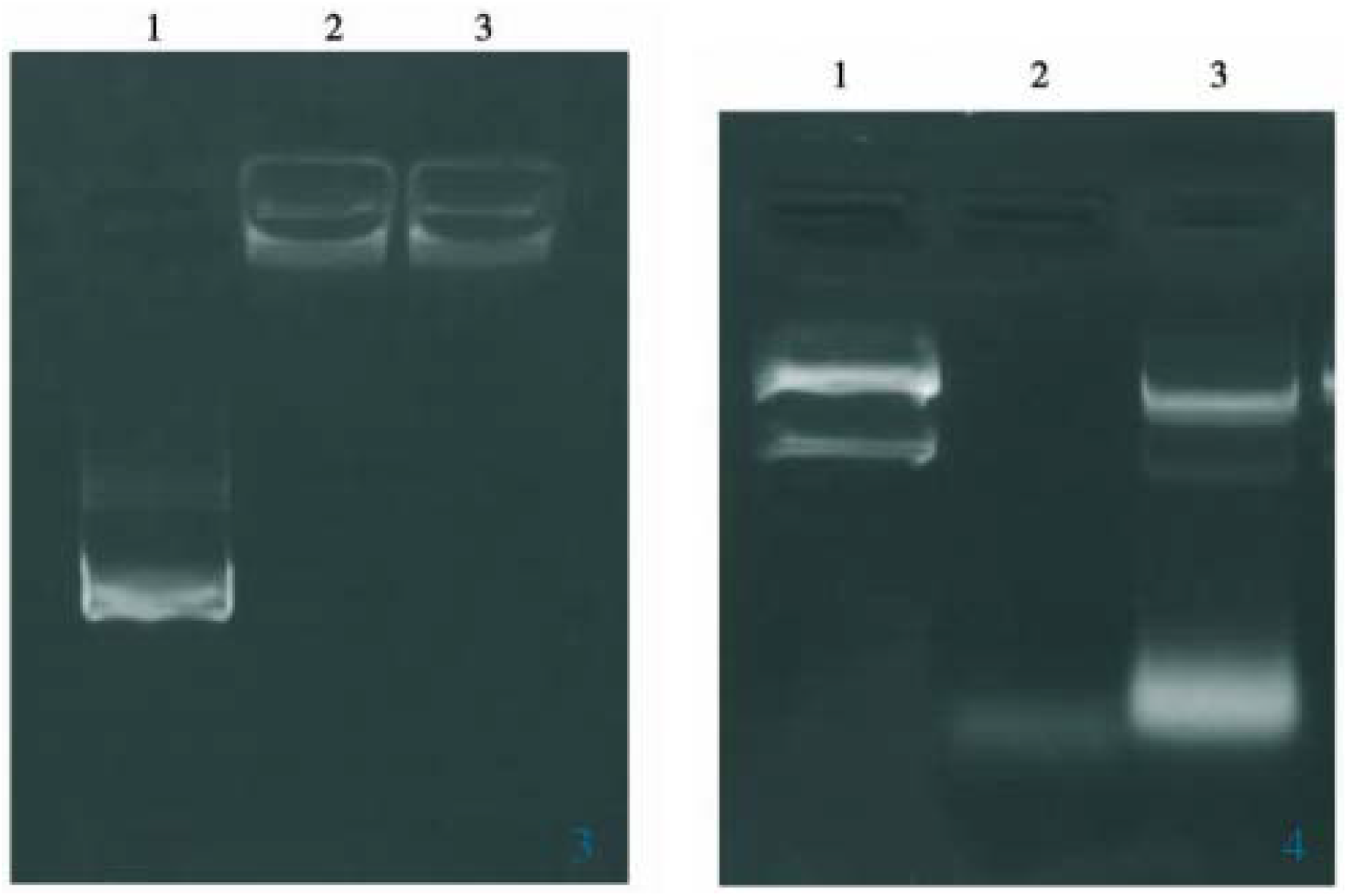
DNA binding efficiency assessed by gel electrophoresis.

The heparin-extracted nanoparticle group was simultaneously subjected to gel electrophoresis with untreated naked plasmid DNA. The results are shown in Figure 4. DNAMePEG-PLGA-NPs remained intact after incubation with DNase I for 30 min at 37 °C. The structure, while the naked DNA was completely degraded, showed that DNA-MePEG-PLGA-NPs can better protect the DNA contained in the DNA from being degraded by DNase I.

**Figure 3.**
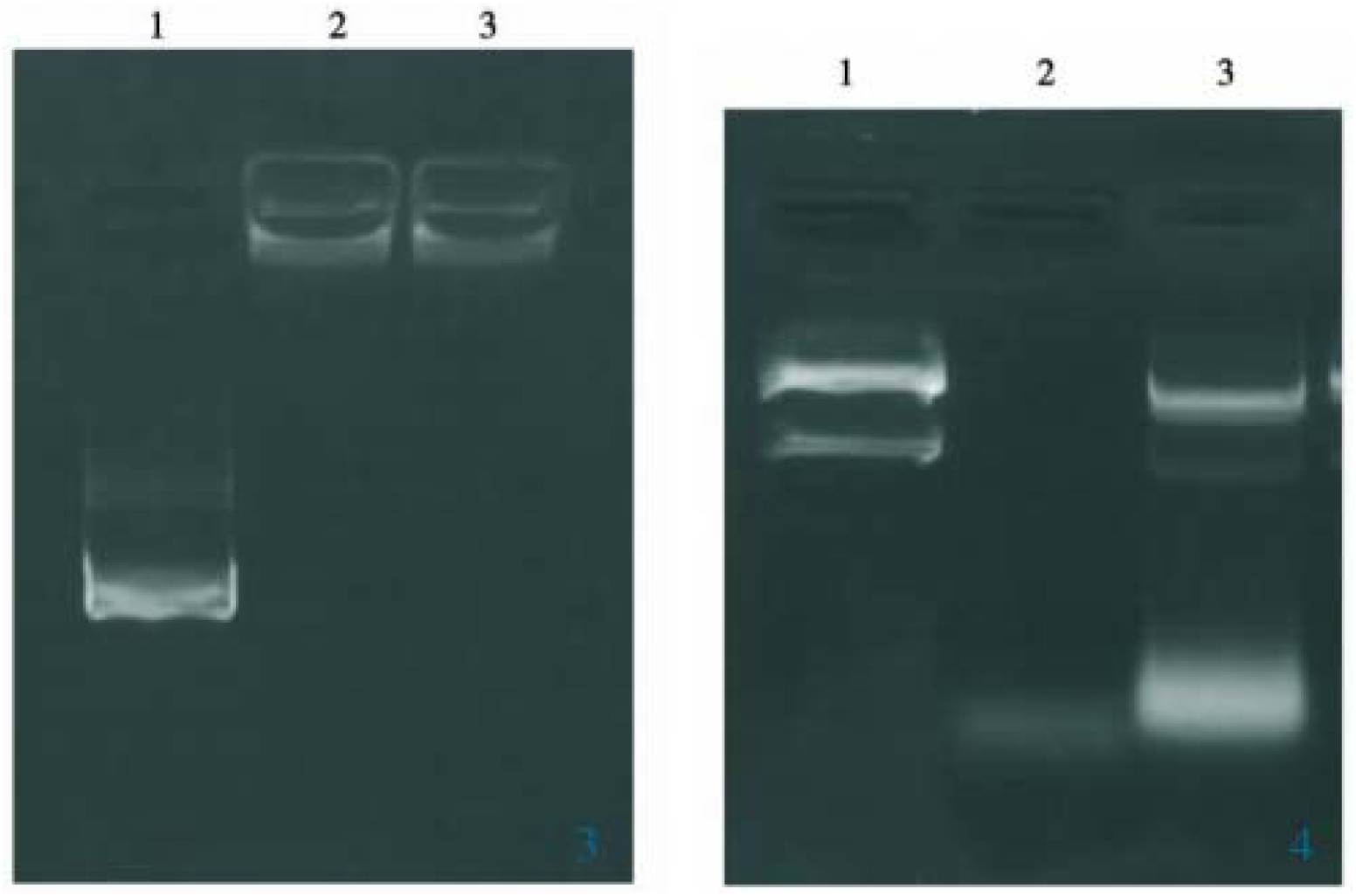
Gel electrophoresis indicate DNA encapsulation efficiency

The cations MePEGPLGA-NPs prepared by the nanoparticle precipitation method are simple in process, controllable in particle size, good in reproducibility, stable in performance, and have certain possibilities to expand production. Acetone is used as the organic solvent alone. Compared with the two organic solvents of acetone and dichloromethane [3], no toxic dioxymethane is introduced, and the interfacial tension between acetone and water is small. When mixed with MePEG – The acetone of PLGA is injected into the aqueous solution of CTAB under stirring. The organic solvent can quickly penetrate the interface, which is conducive to the formation of smaller droplets, and the polymer in the organic phase also migrates to the interface and precipitates, eventually forming smaller nanometers. particle. MePEG-PLGA molecules have a hydrophilic end and a hydrophobic end compared to the nanoparticles prepared by PLGA alone, and MePEG-PLGA-NPs are more stable [16]. The embedding of MePEG changes the softness of PLGA [17], which can form a hydrophilic shell on the surface of the nanoparticle, preventing the agglomeration of the nanoparticles and increasing stability. Moreover, the peripheral MePEG molecule protects the hydroxyl group of PLGA from exposing the surface of the nanoparticle, which greatly increases the surface potential of the nanoparticle compared with the PLGA nanoparticle. The higher the positive potential of the nanoparticle surface, the more capable of adsorbing the negatively charged DNA.

The cationic surfactant CTAB modified MePEGPLGA-NPs has a higher surface potential, allowing the nanoparticles to adsorb more negatively charged plasmid DNA, and the DNA-MePEG-PLGA-NPs surface after adsorption is also higher. The positive charge makes the nanoparticles easy to bind to some glycosyl phosphate on the cell surface, preparing for DNA-MePEG-PLGA-NPs to recognize and enter the cell. The size of the nanoparticles prepared by the nanoparticle precipitation method is close to that of the currently used viral particle carrier, which makes it possible for the nanoparticles to enter the cell by endocytosis. In addition, DNA binding experiments and protection of DNA were not demonstrated by DNase I degradation experiments. The surface scaffold structure of nanoparticles can firmly adsorb and concentrate plasmid DNA, and the DNA is inside the nanoparticle scaffold structure, which is not recognized by nucleases and avoids degradation^12–14^.

The structure of the cationized MePEG-PLGA-NPs combined with the plasmid DNA can be represented by Figure 5. The positively charged nanoparticles form a core-shell structure with the negatively charged plasmid DNA, the nanoparticle is the nucleus, and the DNA adsorbs on the surface of the positive nanoparticles to form a shell. In addition, it can be seen in Figure 1 that DNA-MePEG-PLGA-NPs has a distinct core-shell structure and supports this inference.

In addition, it has been found in many experiments that temperature also has a certain influence on the formation of nanoparticles. When the temperature is high, the shape of the nanoparticles is irregular, the particle size is uneven, and there is adhesion and agglomeration. It is speculated that the temperature may affect the hydrophobic action and volatilization speed of the organic solvent, so the preparation conditions are kept below a certain temperature. The formation of nanoparticles is also necessary.

